# Wild animals suppress the spread of socially-transmitted misinformation

**DOI:** 10.1101/2022.08.23.505022

**Authors:** Ashkaan K. Fahimipour, Michael A. Gil, Maria R. Celis, Gabriel F. Hein, Benjamin T. Martin, Andrew M. Hein

## Abstract

Understanding the mechanisms by which information and misinformation spread through groups of individual actors is essential to the prediction of phenomena ranging from coordinated group behaviours [1–3] to global misinformation epidemics [4–7]. Transmission of information through groups depends on the decision-making strategies individuals use to transform the perceived actions of others into their own behavioural actions [8–10]. Because it is often not possible to directly infer these strategies *in situ*, most studies of behavioural spread in groups assume individuals make decisions by pooling [7, 8, 10, 11] or averaging [8, 9] the actions or behavioural states of neighbours. Whether individuals adopt more sophisticated strategies that exploit socially-transmitted information, while remaining robust to misinformation exposure, is unknown. Here we uncover the impacts of individual decision-making on misinformation spread in natural groups of wild coral reef fish, where misinformation occurs in the form of false alarms that can spread contagiously. Using automated tracking and visual field reconstruction, we infer the precise sequences of socially-transmitted stimuli perceived by each individual during decision-making. Our analysis reveals a novel feature of decision-making essential for controlling misinformation spread: dynamic adjustments in sensitivity to socially-transmitted cues. We find that this property can be achieved by a simple and biologically widespread decision-making circuit. This form of dynamic gain control makes individual behaviour robust to natural fluctuations in misinformation exposure, and radically alters misinformation spread relative to predictions of widely-used models of social contagion.

## Main Text

For humans and other social animals, the actions of others provide constant sources of sensory stimulation that help guide effective decision-making [12–14]. Cues generated by others can encode valuable information about the environment, for example by providing access to stimuli beyond an individual’s own sensory limits [13, 15], or early warning of impending events [15, 16]. But, socially transmitted cues also convey misinformation [4, 5, 14, 17] — erroneous, outdated, or easily misinterpreted content that impedes effective decision-making [17–20]. In stable, long-term groups, animals may solve this problem by inferring the reliability of a signal using knowledge about the sender [12]. However, in many situations such as in crowds [21], automobile traffic [22], bird flocks [1, 20], and animal feeding aggregations [14, 23], decisions must be made rapidly [19, 22] in large or ephemeral [24] collectives, where using foreknowledge of the reliability of all senders is impossible. Behavioural and neurophysiological studies suggest that relatively simple behavioural strategies control decision-making in many such settings [21–23, 25–27]. Yet, whether these strategies somehow account for the possibility of exposure to misinformation is unknown.

To address this question, we deployed camera observatories in a coral reef to continuously record behavioural decision-making of wild, mixed-species groups of foraging fish (Fig. 1; Methods; *Supplementary Information* [*SI*]). Like other social animals [16, 20, 28], reef fish exhibit collective escape responses, wherein individuals within a group cease feeding and flee in rapid succession [23, 24]. Misinformation is produced in the form of escape responses in the absence of true shared threats (i.e., predators), which can spread contagiously through groups as false alarm cascades. In natural foraging groups, escape events in the absence of predators occur regularly, at a mean rate of one event per 7.7 minutes (Fig. 1A). During these events, one individual in the group (the *first responder*) exhibits an escape maneuver [29] involving a deep body bend followed by large acceleration and rapid turning (Fig. 1B; Fig. S1). This initial response may be followed by a cascade of subsequent responses by other individuals (Fig. 1B). Escape behaviour of first responders often coincides with the rapid approach of another fish in the group, indicating an aggressive interaction (Fig. 1D, Fig. S2), to which an escape maneuver is an appropriate reaction. However, aggressive interactions are seldom directed at *secondary responders* (Fig. S2), suggesting that secondary responses are erroneous reactions to a simple form of misinformation: stimuli produced by the aggressor and first responder.

**Figure 1:**
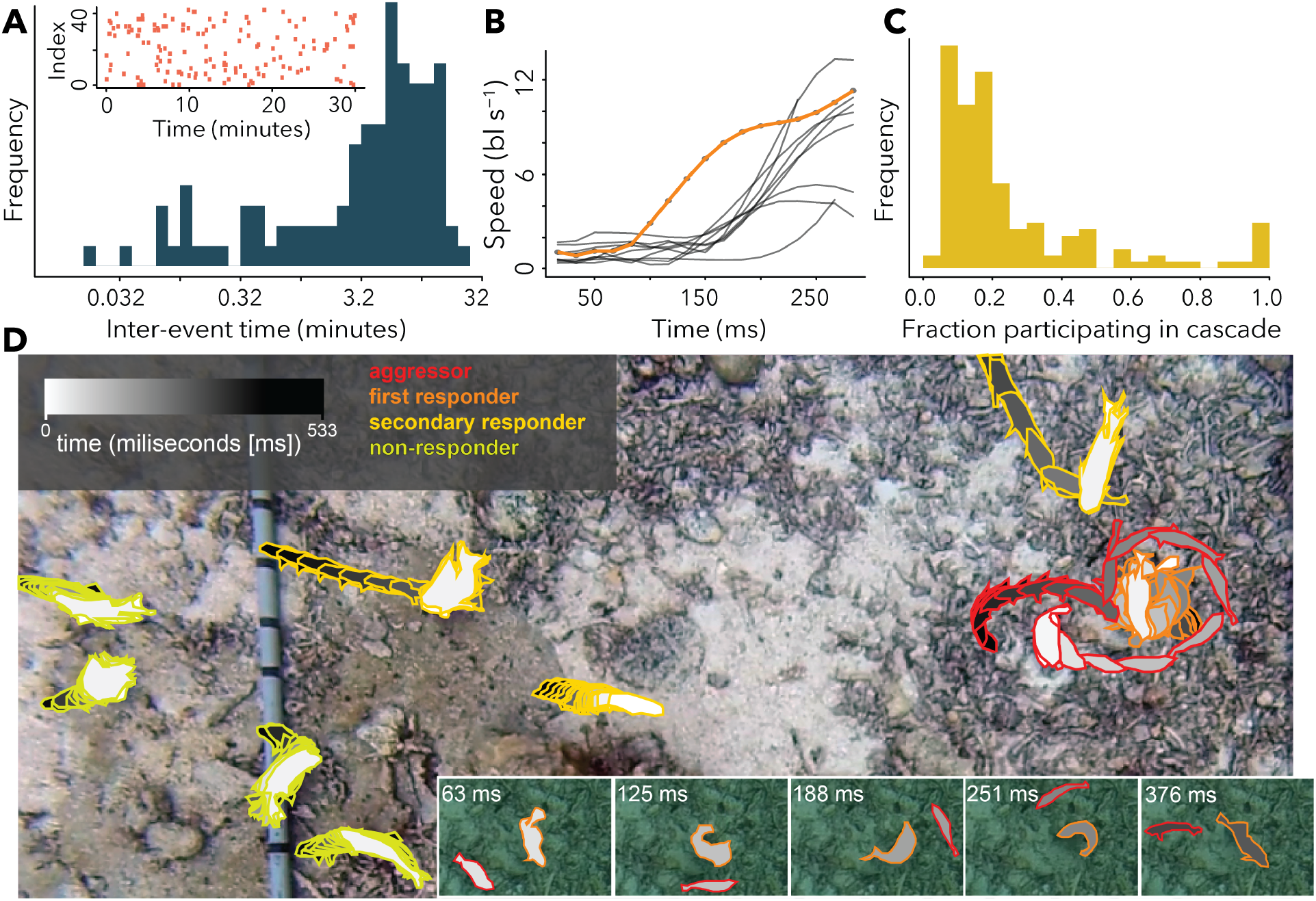
Natural escape cascades. **A.** 42 empirical time series of escape event initiation times (red raster) from observations of unperturbed groups of wild coral reef fish. Histogram shows distribution of inter-event time intervals. **B.** Swimming speed profiles for all responding fish in an example escape cascade (first responder in orange). **C.** Fraction of individuals responding in natural escape events shown in (A). **D.** Example escape event. Aggressor (red) rapidly approaches another individual (first responder, orange), which exhibits an escape response. Secondary responses by other individuals (yellow) follow. Others present do not exhibit escape responses (green). Inset shows time sequence of aggressor and first responder positions.

While some escape events involve large response cascades (Fig. 1B-C), most involve only the first responder. The rarity of large false alarm cascades suggests that individuals employ a decision strategy that is responsive to true threats, like an approaching aggressor, while being robust to the misinformation produced by interactions between other individuals in the group.

In fish, escape responses are controlled by specialized neural circuits that process incoming sensory stimuli, including visual motion stimuli [23], and route it to pre-motor neurons in the hindbrain [30–32]. Experimentally presented visual motion stimuli are sufficient to trigger escape maneuvers of coral reef fish in a manner consistent with known features of these circuits [23]. We therefore hypothesized that natural escape events (Fig 1A-D) are initiated and spread through visual stimuli produced when individuals in the group move. To test this, we reconstructed the visual sensory information available to each animal prior to and during escape events (Fig. 2A; *SI*). As they moved, fish produced strong motion stimuli visible to others (looming motion, translation; *SI*) and these stimuli routinely exceed magnitudes shown in past laboratory experiments to trigger escape behaviour (Fig. S3, [29–32]). Escape responses in our data were preceded by periods of strong looming and translation stimuli from neighbors (Fig. S4); and during escapes, responders turned away from the neighbor producing the strongest stimulus (Fig. 2B). Importantly, individuals that responded also *produced* strong motion stimuli visible to others (Fig. 2C-D), providing a potential mechanism by which the response of one individual could trigger others to respond, thereby propagating a response cascade. To further test this possibility, we developed a simple sensory decision-making model.

**Figure 2:**
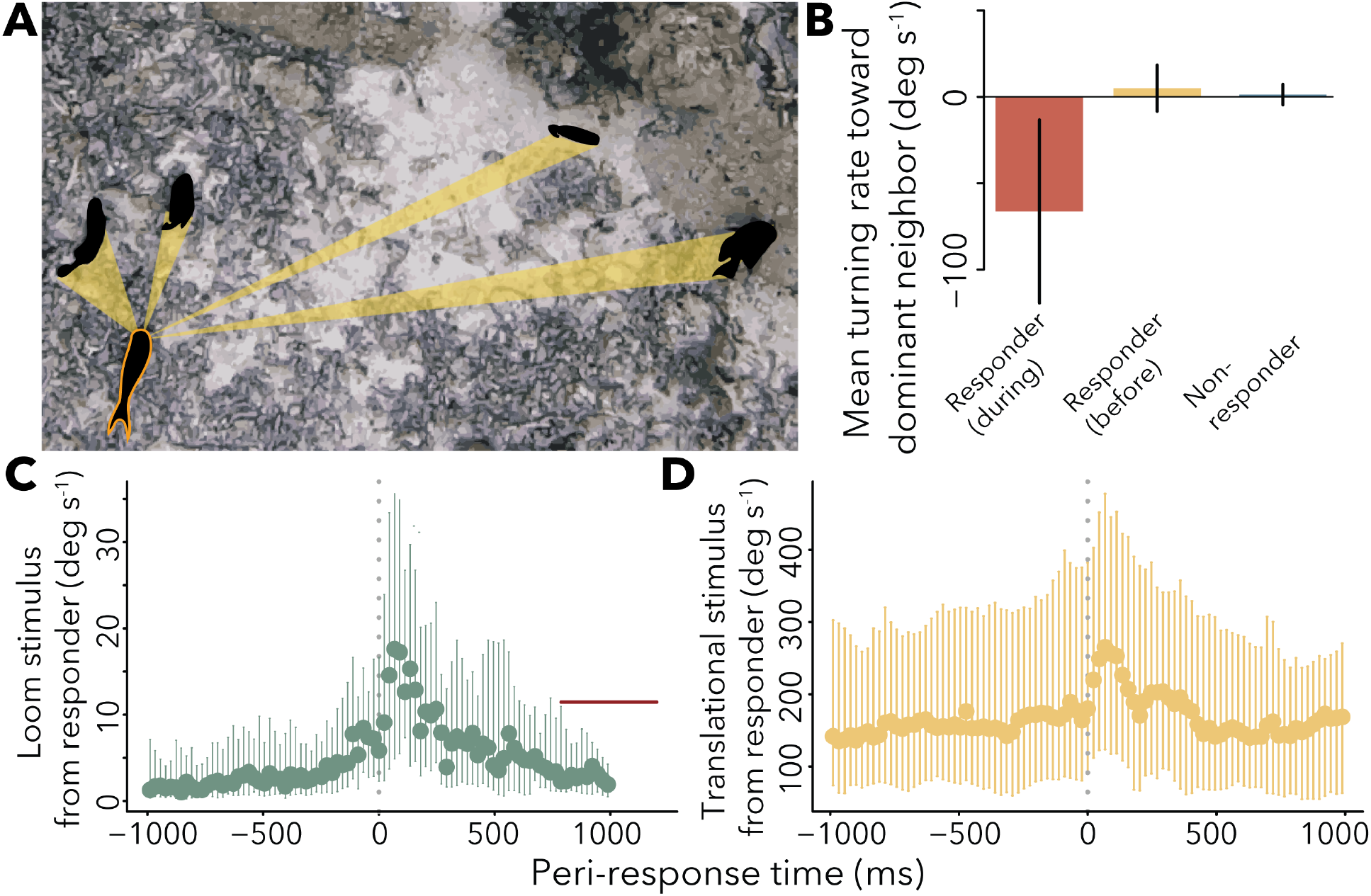
Visual field reconstruction and stimuli during escape events. **A.** All animals were tracked in videos using an automated tracking pipeline (*SI*). A raycasting algorithm [23] and pinhole camera model [26] were used to estimate the projection of each neighbor on retina of focal individual (orange outlined individual). **B.** Mean turning rate toward dominant neighbor (neighbor producing strongest loom) for responding fish within 200 ms of escape initiation time, “responders (during)”; the same fish during periods prior to event, “responders (before)”; and fish present during the escape event that did not respond. Negative values indicate turning away from neighbor. **C.** Median loom rate, and median translation rate (**D**) produced by first responder as perceived by other individuals present. Red line in (C) shows median of putative loom thresholds reported in previous studies [29]. Bars in all panels indicate 25th and 75th percentiles.

Two prevailing hypotheses describe how a decision-maker might operate on sensory information from its neighbors (*SI*, Table S1). Under the first hypothesis, which we refer to as *pooling*, individuals sum sensory input over neighbors [11], respond independently to input from each neighbor [33], or base responses on the strongest input produced by a set of neighbors via selective attention [34]. Under the second hypothesis, which we refer to as *averaging*, individuals average sensory input from different neighbors [25], or average the responses to input from multiple neighbors (response averaging, [26]). To determine whether one of these strategies is consistent with observed escape decision-making, we compared a diverse set of pooling and averaging strategies (Methods; Table S1), and found that no strategy accurately predicted which individuals would respond and which would not (Fig. 3A).

**Figure 3:**
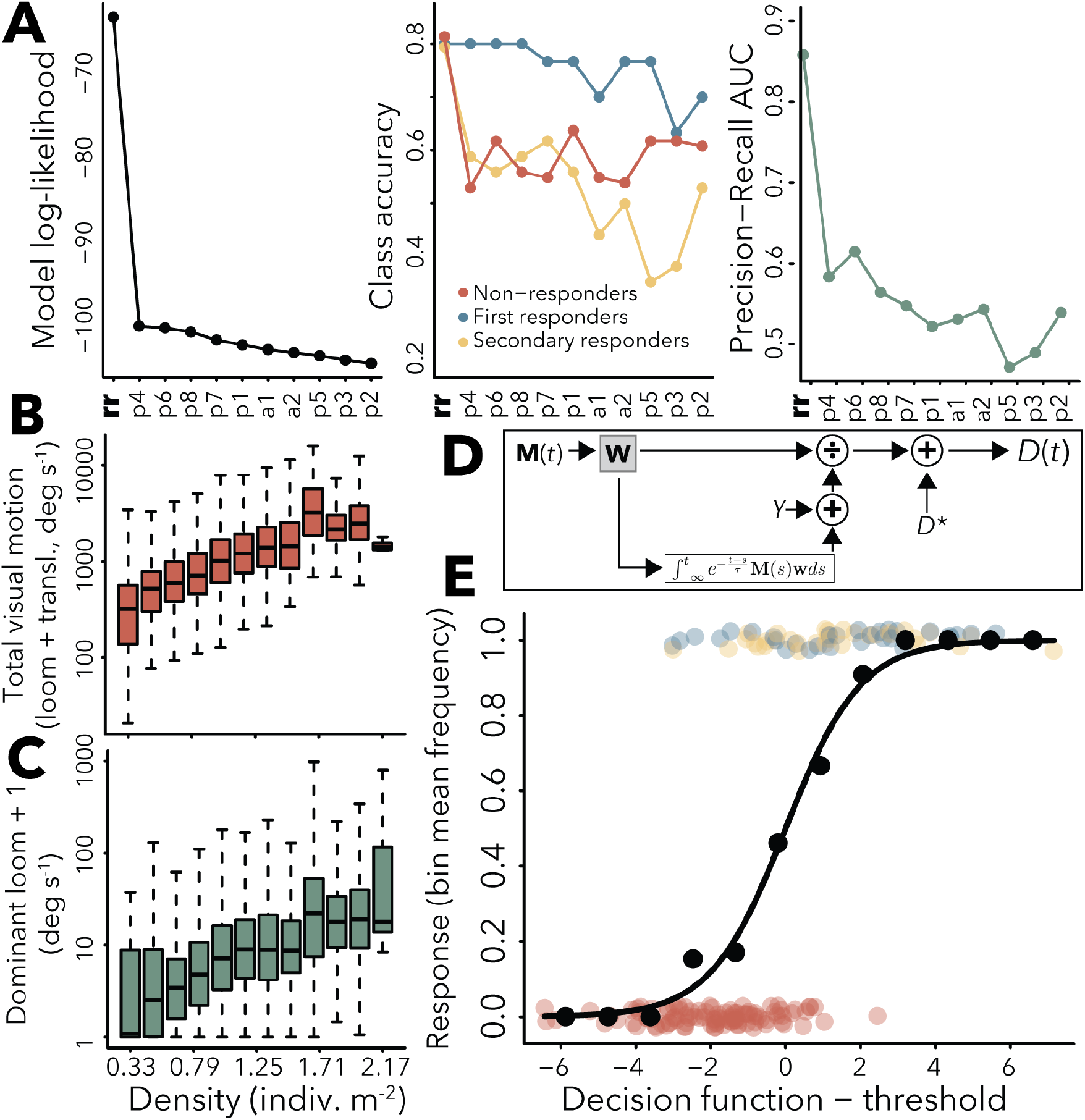
Decision-making models and density scaling of sensory input. **A.** Performance of averaging (“a”), pooling (“p”), and response rescaling (“**rr**”) models: model log-likelihood (left), prediction accuracy for different response classes (center), and precision-recall area under curve (PR AUC, right; see *SI* and Tables S1-S2 for full model descriptions). **B.** Density-scaling of total visual motion input from neighbors, and (**C**) looming from single neighbor producing strongest loom signal. **D.** Diagram of response rescaling model structure (see text, Methods for description of symbols and model derivation). **E.** Observed responses (colored points, colors as in (A) center; points vertically jittered), empirical fraction responding (black points) and predicted (black line) response probability from response rescaling model. Note on (A): we were unable to fit standard phenomenological formulations of simple [7, 8] and fractional contagion [8, 38] models to this data set because, under both models, probability to respond when no neighbor has yet responded is zero; thus these models cannot predict onset of escape events. Nevertheless, we analyze predicted spreading properties of these models following cascade onset in Fig 4.

In reef fish foraging aggregations, the local density of individuals continually fluctuates as individuals enter and leave foraging areas [24]. A curious feature of escape events in our data (Fig. 1A, C) is that, although the total and maximum strength of visual motion stimuli vary by more than 10-fold as local density changes (Fig. 3B-C), escape cascades are typically small (median size = 1 responder; Fig. 1C) and cascade size is uncorrelated with the density of the group during the event (Spearman rank correlation between cascade size and density: *ρ* = 0.093; linear regression, *P* = 0.26 and *R*^2^ = 8.6 × 10^−3^). This suggests that the strategy individuals use to control escape decision-making may involve somehow adjusting sensitivity, or *gain*, applied to sensory input as the overall level of input changes.

Dynamic rescaling of sensitivity is common within sensory organs such as the vertebrate retina [35] that operate across a wide dynamic range of inputs. We asked whether individuals use a behavioural analogue of this process, *response rescaling* [36], to dynamically adjust behavioural responsivity as the volume and intensity of stimuli from neighbors change (see Methods for derivation). One mechanism for achieving such dynamic gain control involves excitation by incoming stimuli along with simultaneous inhibition of responses [37] by the integral of past stimulus input (Fig 3D; Methods). A simple decision rule with this property can be written *D*(*t*) = *D*\* + ***M***(*t*)**w** [*γ* + *m*(*t*)]^−1^, where 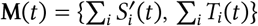 is the vector of perceived looming and translational motion summed over all neighbors, **w** is a vector of constant weights applied to these motion stimuli, *D*\* and *γ* are constants, and 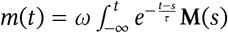**w***ds* is an exponentially-weighted integral of the past history of sensory input from neighbors with constant, *ω*, and decay timescale, *τ*. The probability to respond in some small time increment is given by *p*(*t*) = (1 + *e*^−*D*(*t*)^)^−1^. In this model, sensory input increases the probability to respond, but simultaneously lowers the gain applied to future input, creating an opponency between excitatory and inhibitory effects of visual motion stimuli. In Methods, we show that the dynamic gain control inherent in this model can be achieved by a neural circuit containing just three interlinked populations of neurons. Unlike pooling and averaging strategies, this response rescaling strategy accurately predicts behaviour of first, secondary and non-responders (Fig. 3A, E).

To understand how this decision-making strategy may impact the spread of misinformation, we developed a spatially explicit, empirically-calibrated computational model that simulates populations of individuals, each of which perceives visual stimuli from others and makes decisions to flee or not to flee based on these stimuli (*SI*). In each simulation, we modeled an aggressive interaction between a random pair of nearest neighbors in the population (Fig. 4), and ask how misinformation generated through the interaction travels through the network defined by exchanges of visual information among neighbors. To relate our findings to past work on information spread in social networks [4, 5, 7, 8, 38], we compared populations of individuals that use the response rescaling strategy inferred from our data, to populations in which escape behaviour spreads through the two most widely-used phenomenological models of social contagion: simple contagion [7, 8, 25], under which an individual’s probability to respond is independently influenced by each of its responding neighbors, and fractional contagion [8, 25, 38], under which an individual’s probability to respond depends on the fraction of its neighbors currently responding (*SI*).

**Figure 4:**
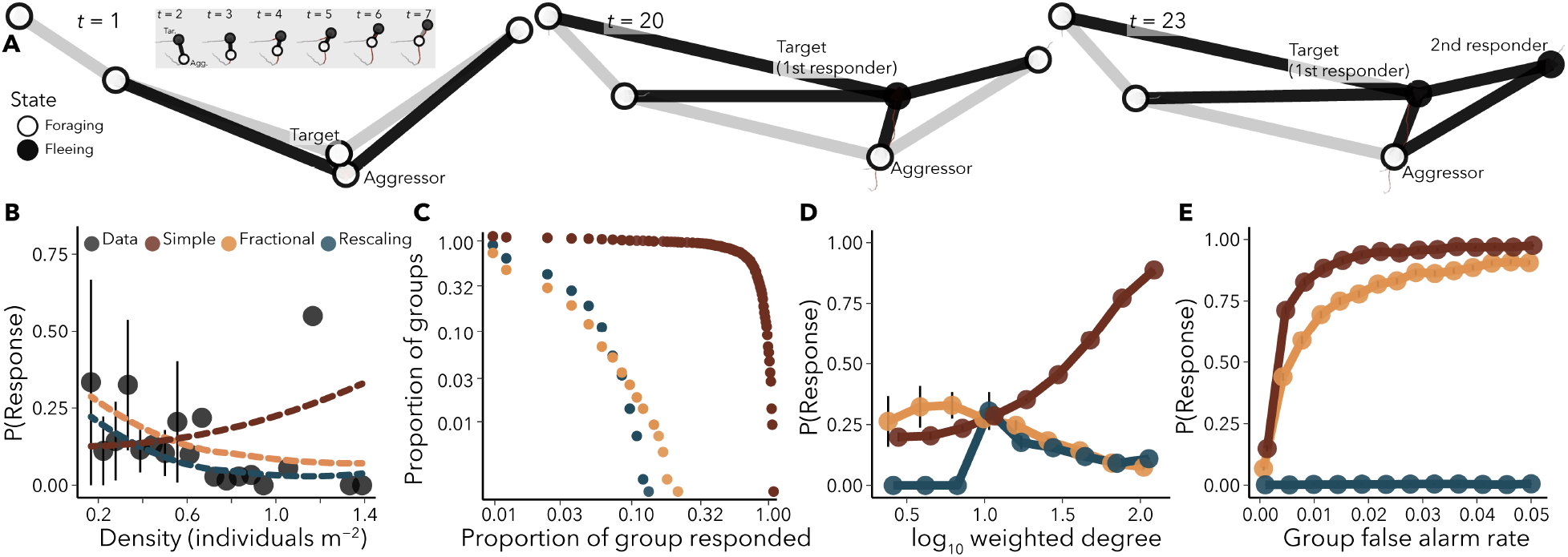
Misinformation spread in populations of decision-makers (computational model). **A.** Time sequence of simulated interaction between aggressor and first responder along with subsequent response of a secondary responder. Edges are drawn between individuals that are within visual range of one another. **B.** Empirical per-capita response probability (points and error bars) as a function of density, and relationships predicted by response rescaling (blue), simple (red) and fractional contagion (yellow) models (*SI*) **C.** One minus the cumulative distribution function of cascade sizes produced by response rescaling, simple contagion, and fractional contagion. **D.** Response probability conditional on exposure to a misinformation cascade as a function of an individual’s weighted degree in the social network. **E.** Probability to respond to misinformation as a function of the per-capita false alarm rate of other individuals in the population.

Consistent with data, the response rescaling population exhibited a decreasing per-capita response probability with increasing density (Fig. 4B) leading to mean cascade sizes that were invariant of density (Fig. S6A). Simulations assuming behavioural spread via simple contagion could not reproduce this pattern, whereas those that assumed fractional contagion produced predictions similar to those of response scaling. The population following fractional contagion also exhibited a similar distribution of cascade sizes to the response rescaling population (Fig. 4C). Fractional contagion shares several spreading properties with response rescaling, in part, because it captures a qualitative property of decision-making in our data: on average, as an individual acquires more neighbors, it requires more sensory input in order to respond (Fig. S5). Thus, rather than acting as superspreaders (Fig. 4D, red curve), highly connected individuals are relatively unlikely to respond to misinformation when they are exposed to it (Fig. 4D, blue and orange curves). This property makes individual decision-making robust to changes in misinformation exposure that occur as density, and as a consequence, local connectivity, changes (Fig. S6B).

Density fluctuations are one source of variation in natural groups [24] that impact exposure to misinformation. Another source is caused by fluctuations in the phenotypic (*i.e.* species) composition of groups, which lead to variation in the rate at which misinformation is generated due to behavioural differences among taxa (Fig. S7). To understand how the probability that an individual will respond to misinformation is influenced by changes in the rate of its generation, we performed simulations in which a focal individual in the population made decisions to escape using either response rescaling, simple contagion, or fractional contagion, and we varied the rate at with other individual in the population generate misinformation by spontaneously fleeing (*SI*, Fig. 4E). The probability that an individual using simple or fractional contagion will respond to misinformation increases rapidly to near one as the origination rate of misinformation increases. In contrast, individuals that use response rescaling maintain a low probability of responding to misinformation that is nearly invariant of the rate at which misinformation is produced (Fig. 4E, blue line). This difference in robustness of decision-making inherits from the dynamic nature of gain control under response rescaling (*SI*): as individuals perceive stronger sensory input from neighbors, inhibitory feedback lowers the gain applied to future visual stimuli from those neighbors, thereby adapting responsivity to the sensory background of the recent past. In this sense, it is motion that is surprising, not motion *per se*, that drives escape responses. Simple and fractional contagion lack this property of temporal adaptation, and decision-making in those models is fragile under changes in the rate of production of misinformation.

Using socially-transmitted information while at the same time avoiding basing decisions on misinformation poses a fundamental conflict for the animal brain. Our results suggest that animals may resolve this conflict by dynamically scaling sensitivity to socially-transmitted cues through simultaneous excitation and inhibition within decision-making circuits. The finding that opponency between social excitation and inhibition is crucial for limiting uncontrolled spread of misinformation may aid in understanding informational processes on human social networks [4, 5, 38], where network architecture and spreading phenomena can be measured, but the rules that govern individual decision-making are not well understood [39].

## Methods

### Data collection

Data were collected in lagoon reefs of Mo’orea French Polynesia. Polyvinyl carbonate (PVC) camera frames (2 m width × 6 m length × 2 m depth) were deployed in lagoon sites on the north shore of the island. Deployment locations were typical of lagoon habitat, and were characterized by a shallow (2.5 m depth) reef flat, comprising primarily pavement and coral rubble adjacent to live and dead colonies of massive and submassive *Porites* corals of 0.5 to 2 m height. Reef fish use these types of open reef flats between coral structure as foraging areas [24]. In each foraging area, a camera frame was mounted using concrete substrate mounts. Foraging areas were filmed continuously from above using downward-facing cameras shooting at either 30 or 60 frames per second (GoPro Hero 3 or Hero 4). Footage used in analyses was collected under unperturbed conditions (i.e., no experimental perturbations were applied), a minimum of 30 minutes after cameras were deployed and researchers had vacated the area. Details on automated animal tracking and visual field reconstruction are described in *Supplementary Information*.

### Comparing models of individual decision-making

The set of plausible models describing how an individual decision-maker might integrate and operate on sensory data from multiple neighbors is vast. We therefore focused on a set of models based on features of decision-making previously described in the perceptual decision-making literature on the basis of behavioural or neurophysiological evidence (Table S1). Multi-source perceptual decision-making models can be organized broadly into two classes on the basis of their assumptions about how stimuli from neighbors are integrated during decision-making: *pooling models* [11, 33, 34], and *averaging models* [2, 25, 28, 40, 41]. Moreover, we consider variants of common decision-making models for each candidate class, including models in which raw visual stimuli are linearly combined to form visual features [11, 33], as well as models in which driving sensory features are nonlinear combinations of raw inputs [23, 42]. Details on models and model-fitting are provided in *SI; Defining the model set* and *Estimating model parameters*.

### Derivation of response rescaling model

We tested a suite of traditional decision-making models and found that none of them accurately predicted individuals’ responses to misinformation (Fig. 3). Based on this result and physiological evidence from other systems [35, 36], we hypothesized that individuals control escape decisions using a rule that dynamically adjusts the gain applied to incoming sensory input based on the recent history of visual input, thereby rescaling responsivity by the sensory background. This type of dynamic gain control is a hallmark of many sensory systems including the vertebrate visual [35] and auditory systems [43], and the chemosensory and internal signalling systems of bacteria and other cells [36, 37, 44], all of which operate over a wide dynamic range of input magnitudes.

Our work considers a distinct but related problem: controlling responsivity to socially-transmitted stimuli amid continual changes in the background levels of those stimuli. We postulated that robust escape decision-making requires two properties: (i) the strength of stimuli necessary to trigger an escape response must vary as the strength of background sensory input changes to allow individuals to maintain sensitivity to changes in visual stimuli when the background level is low, and to avoid becoming overly responsive as the background level increases (Fig. S5); and (ii) the sensitivity of the system should vary as the frequency of events that produce strong sensory stimuli but are not indicative of true threats (i.e., visual misinformation) changes. These properties resemble properties known as Weber’s law — the response of a system to a rapid change in input level is inversely proportional to the background stimulus level; and temporal adaptation – the output of a system after a change to a new background level of input gradually adjusts to a baseline that is independent of the input level [37].

One biologically widespread circuit motif that enables these properties involves fast-timescale excitation by incoming sensory input along with simultaneous inhibition by the same input on a slower timescale [36, 37, 43, 44]. We hypothesize that escape decision-making by coral reef fish is driven by a neural circuit with these properties. In particular we postulate that looming, *S*’(*t*), and translational stimuli, *T*(*t*), are summed over a focal individual’s visual field, such that the driving input for downstream computations is summed visual motion input, 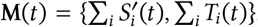. Pooled looming and translation stimuli are then scaled and summed to yield an internal variable, *u*(*t*) = ***M***(*t*)**w**, where 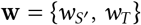 is a vector of constant weights.

The quantity *u*(*t*) could be encoded, for example, in the firing rate of a population of neurons that provides the input to the rescaling circuit. The rescaling circuit involves the firing rate of the input population *u*, the firing rate of a memory [43] population, *m*, and firing rate of a readout population, *y*, with leakage rate 1/*ρ*. The memory population is excited by *u*, and returns to its baseline activity with rate, 1/*τ* in the absence of input from *u*. We take the dynamics of *m*, and *y* to be given by:

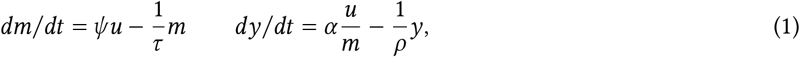

where *ψ* and *α* are constants. Assuming the dynamics of *y* are fast relative to those of *m*, we obtain an expression for the firing rate of the readout population

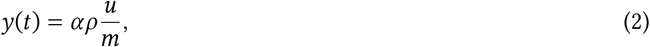

which is proportional to the input, rescaled by the activity of the memory population, *m*. Assuming *t* ≫ 0, yields an expression for the memory population’s activity:

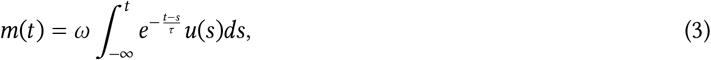

where *ω* = *αρψ*. In practice, for very low activity of the memory population (i.e., *x* → 0), the output of Eq. (2) can become extremely sensitive to fluctuations in input. We therefore introduce a constant, *γ*, to give 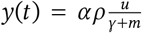. Combining this expression for the activity of the readout population with Eq. (3), and re-writing *y* as the “sensory feature” that drives decision-making (*i.e. F*(*t*) = *y*(*t*)) gives

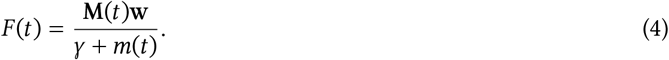

This quantity is assumed to drive decision-making through the function *D*(*t*) = *D*\* + *F* (*t*).

In addition to having the property of exact adaptation and Weber’s law responses to changes in input levels, the model presented in Eqns. (1) exhibits a property known as “fold-change detection”, wherein the entire temporal profile of the circuit’s output in response to a given fold-change in input is invariant of the background input level [37]. This property may be important for maintaining precise control over the timing of signal propagation through the decision circuit irrespective of background input level, which could be crucial given that escape responses must be timed precisely in order to be effective [45]. One candidate module within the fish visual escape circuit [46] that may be capable of performing the computations implied by Eqns. (1) involves populations of glycinergic [47] and dopaminergic [48] interneurons, and tectal projection neurons in the fish hindbrain that receive input from the optic tectum. Although the precise computational properties and connectivity of these populations is still unclear, they have been implicated in a feedforward inhibitory circuit believed to gate escape responses to different types of visual motion stimuli [48].

### Agent-based model details

We designed an individual-based model to simulate the dynamics of populations comprising individuals who follow a response rescaling rule interacting on a featureless 2-dimensional plane. We compared the results of these simulations to expectations from two widely-used models of social contagion: simple and fractional contagion [7, 8, 25, 38, 41]. Model parameters, including those that control agent movement, sensing, and decision-making were set to their corresponding empirical estimates (Table S2). All simulations were performed using the parameter estimates from the best-fitting model. Full details on the individual-based model formulation can be found in *SI*, and an implementation for the Julia 1.7.1 programming language is provided at https://github.com/AshkaanF/scaredyfish.

## Acknowledgements

We thank S. Hein, T. Gross, and S. Munch for comments. A.K.F. was supported by the Research Associateship Program from the National Research Council of the National Academies of Sciences, Engineering, and Mathematics. B.T.M. and A.M.H acknowledge support from National Science Foundation grant no. IOS 1855956.

