## Supplementary Information for "Wild animals suppress the spread of socially-transmitted misinformation"

---

### Initial data Processing

Analyses presented in the *Main Text* focus on natural escape events captured in video footage. Candidate escape events were identified manually by M.A.G. or A.M.H. as times at which one or more individual animals accelerated above typical swimming speeds used during foraging [1]. From this set of candidate events, we designated an event a valid escape event if one or more individual that accelerated exhibited a deep body bend characteristic of a c-start escape maneuver [2] prior to acceleration. All fish exhibiting c-bends followed by acceleration were designated "responders," whereas all individuals not meeting these criteria were designated "non-responders." The first individual to respond in any sequence of responses was designated the "first responder." Due to adverse lighting conditions in many videos, which made automated tracking and visual field reconstruction challenging, only a subset of all collected footage was processed for animal tracking. In tracked videos, designated responses were kinematically similar to experimentally-evoked escape responses from a previous study (Fig. S1, [1]).

### Tracking and Visual Field Reconstruction

#### Video processing and tracking

Prior to tracking, video clips were smoothed using the video stabilization tool in Sony Vegas. Head centroids of all individuals were tracked in two-dimensions using custom detection and tracking software developed in Python using the OpenCV toolkit [1]. All head centroid locations and heading estimates were manually checked by A.M.H. or M.A.G., and locations and heading estimates were manually corrected if necessary using a track-correction application developed by G.F.H. After tracking, position and heading estimates from the subset of videos that were recorded at 30 frames per second were upsampled to 60 frames per second using linear interpolation, yielding a constant sampling rate of 60 position and heading estimates per second for all individuals in all tracked clips. All trajectories and headings were smoothed using a $5th$ order Savitzky-Golay filter with a filter length of 0.183 seconds (11 frames at 60 fps). Finally, the body length of each individual was computed by measuring the distance (in units of pixels) from the snout to tail tip. All length estimates were converted to metric units using physical waypoint markers on the base of camera frames visible in all video clips.

#### Pinhole camera model

For each individual in each tracked clip (henceforth "focal fish"), we reconstructed the perceived visual sizes and locations of all other fish (henceforth "neighbors") at every point in time using a "pinhole camera model" similar to that used by Hein *et al.* [1] and Harpaz *et al.* [3]. For each focal fish at each time point, we registered the head locations, snout locations and tail tip locations of all neighbors in the egocentric reference frame of the focal fish with the focal fish's head centroid placed at the origin. Rays were then cast from the origin in the horizontal plane at a resolution of $5 \times 10^{-3}$ radians per ray, and the number of rays intersecting the line segment connecting the snout and tail tip of each neighbor was used to compute the angular width of that neighbor from the perspective of the focal individual. For each ray, we also estimated the visible area in the vertical (i.e. depth) dimension assuming the body height of each neighbor was one fifth its length. We used this measure as an estimate of the angular height of the neighbor from the perspective of the focal fish. We defined the angular size, $S$, of each neighbor as the geometric mean of its angular height and angular width.

### Formulating Models of Decision-Making During Escape Events

#### Relevant visual sensory inputs

We constructed a set of decison-making models based on previously studied properties of the fish visual escape circuit, and visually-evoked escape behavior in fishes and other animals [1, 2, 4–9, 9, 10]. Based on this work, we restricted our analysis to three classes of visual stimuli produced by neighbors: the angular size of a neighbor, $S$ [4, 5], defined as the geometric mean of the angular length and height of each neighbor computed from the pinhole camera model (see *Pinhole camera model* above), looming motion [4, 5, 7, 8], $S'$ defined as $\max(0, dS/dt)$, where $dS/dt$ is the time-derivative of angular size estimated through finite differences of smoothed angular size, and translational motion [8], $T$, defined as the self motion-corrected rate of change in angle from the head centroid of the viewer to the head centroid of the neighbor (see *Correcting translation for self-motion*), which encodes motion of a neighbor in the plane orthogonal to the focal individual's line of sight to that neighbor. Loom rates were set to a minimum of zero based on published experimental results showing weak effects of receeding (i.e., $dS/dt < 0$) on escape responses of fishes [4].

Effects of stimulus loom rate and angular size have been analyzed in many studies of visually-evoked escape behavior in fishes [1, 2, 4, 5, 8], and both variables have been invoked as possible sensory stimuli responsible for triggering responses [2, 4, 5]. In contrast, few studies have explored effects of translating stimuli on escape responses; however, theory suggests that translational motion could be important for avoiding collisions [11], and a recent study by Förster *et al.* [8] found that some classes of looming and translating visual stimuli can elicit patterns of neural activity in the optic tectum that overlap significantly, potentially implicating translational motion as a stimulus capable of influencing escape responses.

**Correcting translation for self-motion.** Self-motion is known to create strong optic flow (*i.e.* whole-field translational motion) through the apparent relative motion of stationary objects in an animal's visual field as that animal moves. Because of this, turning or forward-backward motion of a focal individual can create strong apparent translational motion stimuli from neighbors. From the perspective of the focal animal, mechanisms of gaze stabilization [12] and efference copy modulation [13, 14] can compensate for this self motion-induced translation, allowing an animal to discriminate externally generated, and self-generated translational motion.

To account for such biological mechanisms of self-motion correction, we compute the translation signal used in models of decision-making in the following way. For a focal individual, we first computed raw translational motion of the $i$th neighbor at time, $t$, by finite differences as follows: $T_{\mathrm{raw},i}(t) = [\theta(\mathbf{y}(t), \mathbf{x}_i(t)) - \phi_i(\mathbf{y}(t-\delta), \mathbf{x}_i(t-\delta))]\,\delta^{-1}$, where $\phi(\mathbf{y}(t), \mathbf{x}_i(t))$ is the angular location of a neighbor located at two-dimensional position $\mathbf{x}_i$ in the visual field of a focal individual located at position $\mathbf{y}_i$, and $\delta$ is the time increment. We then computed the deviation of observed translation of the $i$th neighbor from the optical flow cue produced if the neighbor were stationary and, therefore, all raw translation were generated solely by self-motion: $T_i(t) = T_{\mathrm{raw},i}(t) - [\phi(\mathbf{y}(t), \mathbf{x}_i(t-\delta)) - \phi_i(\mathbf{y}(t-\delta), \mathbf{x}_i(t-\delta))]\,\delta^{-1}$. The latter term in brackets represents the expected optical flow from the *ith* neighbor if it were stationary. Substituting the expression for $T_{\mathrm{raw},i}(t)$ gives $T_i(t) = [\phi(\mathbf{y}(t), \mathbf{x}_i(t)) - \phi(\mathbf{y}(t), \mathbf{x}_i(t-\delta))]\,\delta^{-1}$.

### Defining the model set

To convert a stream of continuous visual inputs (i.e. the size, loom rate, and translation rate from each neighbor) into a binary decision to respond or not to respond, each decision-making model requires three elements: (1) a function that transforms the relevant raw sensory variables into latent "sensory features" [15] on which decision-making is based, (2) a description of how incoming sensory input or sensory features from each neighbor are combined [16], and (3) an output function that maps the values of these sensory features to a response probability, $p$. Few models in the literature fully specify all three of these elements. One reason for this is that many perceptual decision-making models were originally developed to describe decision-making in response to stimuli from a single source (e.g., [9, 17–19]), and therefore do not specify how stimuli from elsewhere in the visual field are integrated. In the case of several other models [16, 20, 21], the driving input from each neighbor is taken to be the neighbor's "behavioral states" (0 or 1 indicating whether the neighbor is or is not engaging in the behavior of interest, e.g., [16]) or a random draw from a statistical distribution with parameters that are functions of the neighbor's state (e.g., [22]) rather than the values of sensory stimuli perceived by the decision-maker. To develop a set of alternative models to apply to our data, we organized and adapted previously proposed models in a way that allowed us to accommodate our data.

**Pooling models.** We considered three classes of *pooling models*: (1) models that assume the decision-maker sums sensory inputs over neighbors and bases decisions on this summed quantity [23], (2) models that assume the decision-maker operates independently on sensory input from each neighbor and changes its behavior when input from at least one neighbor is sufficient to trigger a response [24], and (3) models often referred to as "winner-take-all selective attention" models, which assume the decision-maker bases decisions only on sensory input from the neighbor producing the strongest sensory stimuli at a given time [7]. Mathematical descriptions of these models are given in Table S1.

In addition to models in which raw visual stimuli are linearly combined to form visual features, we consider several models in which driving sensory features are nonlinear functions of raw inputs. In particular, Hatsopoulos *et al.* proposed that escape decisions in response to looming from a single source may be driven by a nonlinear visual feature of the form $F(t) = S'(t)\exp[-\beta_2 S(t)]$ [17], where $\beta_2$ is a constant. Hein *et al.* [1] performed a multi-model comparison and identified a modified form of this model that well-described data, which in the present context can be approximated as $F(t) = S'(t)\exp(-\sum_i \beta_2 S_i(t))$, where $i$ indexes neighbors of the focal decision-maker and $\beta_2$ is a constant. In classic work, Lee [19] observed that the time until collision with a looming object moving at constant speed could be inferred by a sensory feature given by the proportionality $F(t) \propto \frac{S(t)}{S'(t)}$. Finally, several studies [4, 5, 25] have proposed heuristic models positing that responses to looming objects occur when the visual size of a looming object crosses a threshold. A model with this property can be formalized by making the probability to respond a function of the sensory feature, $F(t) = \beta_1 \mathbf{1}_{[S'(t)>0]} S(t)$, where $\beta_1$ is a constant and $\mathbf{1}_{[S'(t)>0]}$ is an indicator function that takes a value of one if the neighbor is looming and zero if it is receding (Table S1).

Models of winner-take-all selective attention (i.e. decision-maker responds only to a single dominant stimulus source at any given time) require designation of which neighbor is dominant. For each focal individual at each time, we designated the dominant neighbor as the visible neighbor producing the strongest loom signal, based on the finding from past studies that dark looming elicits stronger and more consistent neural and behavioral responses than do other visual stimuli [4, 8]. In cases in which no neighbor was looming (i.e., all were receding), we designated the neighbor producing the strongest translational motion stimulus as the dominant.

**Averaging models.** In addition to pooling models, we considered *averaging models*. One major class of averaging model proposed in the literature treats sensory input from each neighbor as binary (one indicates that the neighbor is engaging in the behavior of interest and zero indicates that it is not) and computes whether the focal decision-maker will change state as a function of the average (e.g., [16, 21]) or sensory-weighted average (e.g., [22]) of the binary states of visible neighbors [16, 20–22, 26]. In the present study, we model sensory input from neighbors as continuous variables (i.e., rate of looming motion, translation rate, visual size) rather than binary states, based on evidence from neurophysiological studies showing that the visual system encodes continuous variation in visual size and motion stimuli [4–8] and that stimulus magnitude is important in driving responses [1, 7, 25]. In this context, a model analogous to those studied by Strandburg-Peshkin *et al.* [21], Rosenthal *et al.* [22], Sosna *et al.* [26], and Poel *et al.* [16, 20] treats the probability that a decision-maker will change state at time, $t$, as a function of the average continuous sensory input computed over all visible neighbors (model a1 in Table S1).

A second class of averaging model assumes that the behavioral response to inputs from multiple neighbors is an average of the behavioral responses to each individual neighbor [3]. Note that this is also the effective assumption underlying many so-called "flocking models" of collective motion (e.g., [27, 28]). In our setting, behavioral responses are binary (i.e., an individual either initiates an escape response or does not initiate a response), and, therefore, a model in which responses are averaged cannot be applied directly. However, a model analogous to that of Harpaz *et al.* [3] can be constructed by postulating that the probability to respond, $p$, in some small increment of time, is given by $p = \frac{1}{N}\sum_i p_i$, where $p_i$ is the probability to respond to the $i$th visible neighbor if it were the only neighbor present, and $N$ is the number of visible neighbors (model a2 in Table S1).

**Response rescaling.** The response rescaling hypothesis proposed in the Main Text postulates that individuals may control escape decisions using a decision rule that dynamically adjusts the gain applied to incoming sensory input based on the recent history of visual input, thereby rescaling responsivity by the sensory background. A derivation of this model can be found in the *Methods* section of the Main Text.

### Comparing Decision-Making Models to Data

#### Estimating model parameters

To compare predictions of decision-making models to escape decisions exhibited by animals in our data, we began by computing time series of the three visual stimulus variables — neighbor visual size, neighbor looming, and neighbor translation — for all individuals present in all tracked escape events. We truncated the time series for each individual to begin one second prior to the onset of the first responder's escape response, and to end at either (i) the onset time of that individual's escape response (if the focal individual responded), or (ii) one second after initiation of the last escape response in the escape event (if the focal individual did not respond). Aggressors (i.e. the individual that charged another individual in trials where this behavior was evident) were excluded from analysis. We used each individual's time series of visual inputs to compute a time series of sensory feature values $F(t)$ (Table S1). We then averaged the value of this sensory feature over the time series to provide an estimate of the overall level of sensory stimulation experienced by each individual, $\overline{F}$. We used model equations given in Table S1 with $\overline{F}$ as the sensory feature value to generate a response probability prediction for each individual in the dataset.

For each model in the model set, we estimated parameters by minimizing the negative log-likelihood of the data conditional on parameters over the entire set of tracked individuals (i.e., all responders and non-responders). The amounts to minimizing

$$NLL = -\sum_k \log[\text{prob.}(z_k|\theta)], \tag{1}$$

where prob.$(\cdot|\theta)$ is the probability that the $k$th individual will exhibit response $z_k$ (response or no response), conditional on the model and parameters $\theta$.

With the exception of two models (models p2 and a2 in Table S1), the response probability for the $k$th individual in each model is assumed to follow the relation

$$\text{logit}(p_k[t]) = D^{\star} + F_k(t - \Delta), \tag{2}$$

where $\text{logit}(x) = \ln\left[\frac{x}{1-x}\right]$ is the logit function, and $\Delta$ is a sensory-motor delay. The sensory-motor delay accounts for the time delay between the arrival of visual stimuli at the retina and the time at which the motor system can begin executing a command in response to those stimuli [18]. In all models, we assumed this delay time was 30 milliseconds based on a prior analysis of coral reef fish escape behavior in response to visual stimuli [1].

In the case of the two models in which the relation in Equation (2) did not hold – model $p_2$ and model $a_2$ (Table S1) – the expression for response probability is given in Table S1. Because our data (Fig. S1) and a previous study [1] documented differences in responsivity among species in different families, we allowed the logit intercept parameter, $D^{\star}$ in all models to differ depending on the family to which each individual belonged. Based on past work [1], the family groupings used were: Acanthuridae, Scaridae/Labridae, Zanclidae, and a single group containing individuals in all other families observed in the data.

For models that had the form shown in Equation (2), we fitted all linear terms using the `glm` function in R, gridding over nonlinear parameters if necessary. For models that could not be cast as a generalized linear model (i.e., models p2, and a2, Table S1), we instead performed a brute force grid search to identify optimal values of all parameters. For each parameter set, we computed the negative log-likelihood, and we compared this value among all parameter sets to identify the parameter values that minimized the negative log-likelihood.

#### Evaluating model performance

Model performance measures for all fitted decision-making models are provided in Table S2, and in Figure 3A in the Main Text. We report three measures of model performance. Firstly, the negative log-likelihood was computed for each model following Equation (1). As a second measure of model performance, we computed the area under the precision-recall curve (PR AUC) for each model. This statistic quantifies how the model trades off true positives and false negatives, which provides a measure of the tradeoff between the precision of predicted responses (*i.e.*, when the model predicts that an individual will respond, how likely is that prediction to be accurate?) and the fraction of true responses in the data that are correctly predicted. We computed PR ACU by computing precision (precision $= n_{tp}[n_{tp} + n_{fp}]^{-1}$, where $n_{tp}$ and $n_{fp}$ are the number of true and false positives predicted by the model, respectively) and recall (recall $= n_{tp}[n_{tp} + n_{fn}]^{-1}$, where $n_{fn}$ is the number of false negatives predicted by the model) for each of $n = 1000$ threshold values, spaced evenly between zero and one. We then numerically integrated the precision-recall curve using a trapezoidal integration scheme. Finally, we computed model prediction accuracy for each class of individuals in the data: first responders, secondary responders, and non-responders, where the prediction accuracy for each class is the fraction of observations in each class predicted correctly by the model. To convert predicted response probabilities into binary response predictions (i.e. escape response or no escape response), we had to specify a response threshold for each model, such that predicted probabilities above that threshold were predicted as responses and those below the threshold were predicted as non-responses. Because we were interested in identifying models that could correctly predict responses for all classes, we computed a threshold for each model by computing the prediction accuracy for each class using $n = 1000$ threshold values, spaced evenly between zero and one. For each threshold value, we then computed the standard deviation of accuracy across the three classes and selected the threshold that minimized this quantity. Note that neither negative log-Likelihood nor PR AUC require specifying a threshold, and thus the ranking of models with respect to those metrics is independent of this choice.

### Individual-based Model Formulation

To understand the dynamics of populations comprising interlinked individuals who follow a response rescaling rule, we developed an individual-based model that captures the key elements of the sensing and decision-making algorithms inferred from our data. An implementation of the model in the Julia 1.7.1 programming language is provided at https://github.com/AshkaanF/scaredyfish.

Model parameters, including those that control agent movement, turning, sensing, and decision-making were set to their corresponding empirical values or averages where estimates from our data were possible (Table S3). In addition to these empirically-determined parameter values, we calibrated two free parameters — $D^\star$ and $w_{s'}$ — using a grid search procedure, to minimize the root-mean-square error between simulated agent response probabilities and empirical response probabilities across the empirically-observed range of fish densities (Fig. 4B). Based on parameter estimates from the best-fitting response rescaling model to data (Table S2), we set $w_T = 0.21 w_{s'}$. All subsequent simulations were performed using the best-fitting parameter set.

#### Agent morphology and movement

Individuals in the model are represented as 20cm × 4cm (diameter × height) cylinders, dimensions equal to the mean empirical body sizes of reef fish in our study, moving on a featureless two-dimensional plane. We assume that each agent has a 360° field of vision that it uses to sense other nearby individuals within a fixed visual range, $\kappa$. Depending on the perceived visual motion of neighbors (see *Relevant visual sensory inputs*), agents switch between two states: a *foraging* state, in which individuals follow a random walk and movement is relatively slow, with speeds that match those of real animals during foraging bouts; and a *flee* state, in which individuals accelerate up to a speed of 7 body lengths $s^{-1}$ while steering away from neighbors using the same directional bias as real fleeing fish (Fig. 2B; Table S3). In the foraging state, each agent $i$ follows a correlated random walk by adjusting its current heading $h_{i,\text{curr}}$ according to $h_{i,\mathrm{curr}} + N$, where $N \sim \mathcal{N}(0, u_0)$, and taking a step of size $W$, which is drawn independently from $W \sim \mathcal{U}(0, w_0)$ in each time step (see Table S3 for details on parameters).

#### Agent sensing and decision-making

Agent sensing and decision-making was designed to mimic the visual motion-based rules inferred from field data. We therefore encoded the visual system of each agent as a two-dimensional visual field using the same pinhole camera model as in visual field reconstructions for real animals (see *Pinhole camera model*). In each time step, agents measure the loom rates, self-motion-corrected translation rates, and angular sizes of all visible neighbors (see *Relevant visual sensory inputs* for calculations). Because agents in our model are morphologically identical and rigid, partial and whole visual occlusion of neighbors was computed using a straightforward occlusion tree-like algorithm, as opposed to a discrete ray-casting method. To avoid numerical artefacts arising from the interaction between visual occlusion and finite difference estimates of motion features, we assumed that a semi-occluded neighbor would be discernible to a focal individual if at least 10% of its body was visible.

During simulations, the probability that an agent will switch from the *foraging* to the *flee* state in any given time step depends on both the current and recent history of observed looming and translational motion according to Eq. 4, Main Text (Table S1). Thus, in addition to tracking the current (*i.e.* time-lagged) perceived motion of all neighbors, agents retain a 0.6-second finite working memory of these motion features, which is used to calculate the activity of the inhibitory memory population, $m$ (Eq. 3, Main Text). The decay timescale of memory $\tau$ was set to 0.1 seconds for all simulations based on estimates from the best-fitting model to data (Table S2). Flights were assumed to last for 0.69 seconds, which reflected mean observed empirical flight durations.

#### Numerical experiment for misinformation spread

Two main numerical experiments were performed to understand the dynamics of simulated populations comprising individuals following the response rescaling rule. The first set of simulations had the goal of understanding how a single unit of misinformation travels through networks of interlinked individuals, where the undirected links of the network mean that two individuals are within each others' sensory radii and are not visually occluded. To mirror the natural process seen in the field, we define a unit of misinformation as a pairwise antagonistic interaction between an aggressor and a target (Fig 4A, Main Text). To simulate this interaction, we initialize each instance by randomly placing individuals in a square plane with dimensions that achieve a desired initial density, and then randomly choosing an agent to be the aggressor. The aggressor follows a pure pursuit [29] strategy as it closes the distance to its nearest neighbor. The aggressor ceases its charge once a minimum distance to the target neighbor is achieved; only simulations in which the aggressor successfully completes a charge, and the target responds by switching to the *flee* state were retained for analysis. Initial agent positions and headings for simulations are chosen randomly to achieve desired group densities, while maintaining a constant number of agents. All simulations were performed with $N = 100$ agents, although our results were insensitive to the number (as opposed to the density) of agents.

To understand how variation in the frequency of exposure to misinformation would impact decision-making in individuals who deploy response rescaling, we performed a second set of computational simulations holding density at its empirical average, $\hat{N} \sim 0.8$ individuals m$^{-2}$. Here, a single focal individual in a population of $N = 100$ individuals is given decision-making capabilities (*i.e.* the response rescaling rule), while all other individuals spontaneously switch to the *flee* state with a constant probability $p$ in each time step. Results of these simulations are summarized as plots showing the relationship between the rate of spontaneous startles $p$ and the response probability of the decision-maker (Fig. 4E).

#### Comparisons to phenomenological models of social contagia

We sought to compare the results of our numerical experiments with predictions from two widely-studied models of social contagia: simple and fractional contagion [16, 30, 31]. In these alternative models, agent movement rules and visual capabilities are the same as described above (Table S3), with identical parameters and numerical experiments in all cases. The difference in these alternative models is in the decision function, in that agents no longer use a response rescaling rule (Eq. 4 Main Text) to decide when to switch to the *flee* state. Instead, for the case of a simple contagion model, response probabilities depend on the state of the focal agent's neighbors according to $P(\text{Response}) = 1 - (1-q)^{n_{\text{flee}}}$, where $n_{\text{flee}}$ is the number of visible neighbors in the *flee* state, and $q$ is a parameter that is calibrated to empirical cascade size distributions using the grid search procedure described above. Responses in the fractional contagion model are described by $P(\text{Response}) = \frac{1}{n}\left[1-(1-q)^{n_{\text{flee}}}\right]$, where $n$ is the number of visible neighbors [16, 20, 22, 30]. As was done for the response rescaling model, we calibrated both simple contagion and fractional contagion models to minimize the root-mean-square-error between empirical and simulated false alarm cascade distributions across empirically-observed fish densities (Fig. 4B, Main Text). Here, we tuned only a single parameter, $q$, which was estimated to be $q = 0.00383$ for the case of simple contagion, and $q = 0.0213$ for fractional contagion.

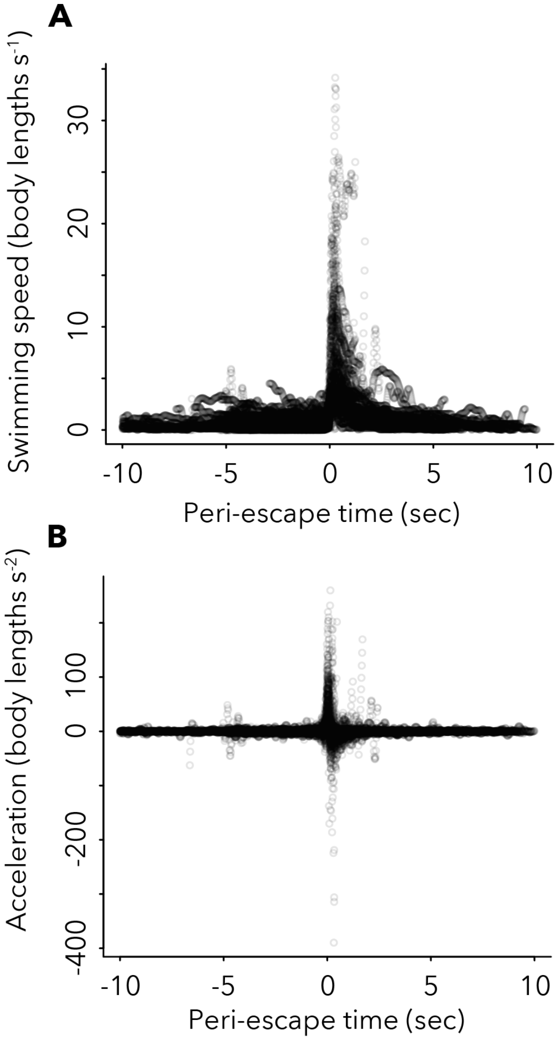


**Figure S1. Kinematics of escape responses. A.** Swimming speeds and **B.** accelerations of fish designated as "responders" during escape response events. Time axis shows time relative to the frame at which the body bend [32] at the onset of escape responses began (see SI section *Data Collection and Processing*).

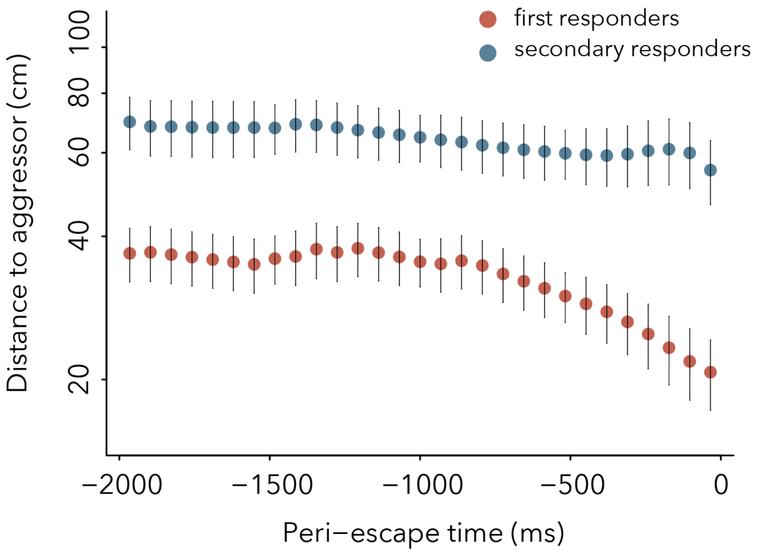


**Figure S2. Distance to aggressor.** Mean distance between aggressor and responders during period leading up to response onset for trials in which there was a clear aggressive interaction. Distances are shorter for first (red) than secondary responders (blue), and distance between aggressor and first responders shrinks rapidly prior to onset of escape response for first responders but not for secondary responders.

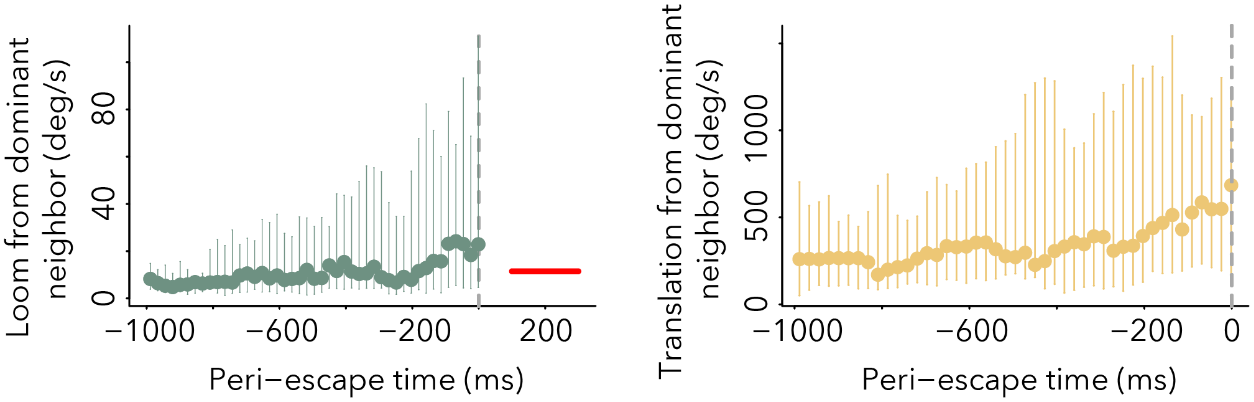


**Figure S3. Visual stimuli to first responder.** Instantaneous loom rate, $S'(t)$, (left) and translation rate, $T(t)$ (right) from dominant neighbor (neighbor producing the strongest looming stimulus) perceived by first responder during period preceding initiation of escape response (at time 0). Bars indicate 25th and 75th percentiles. Red line shows median of putative lower loom thresholds reported in previous studies of fishes [32].

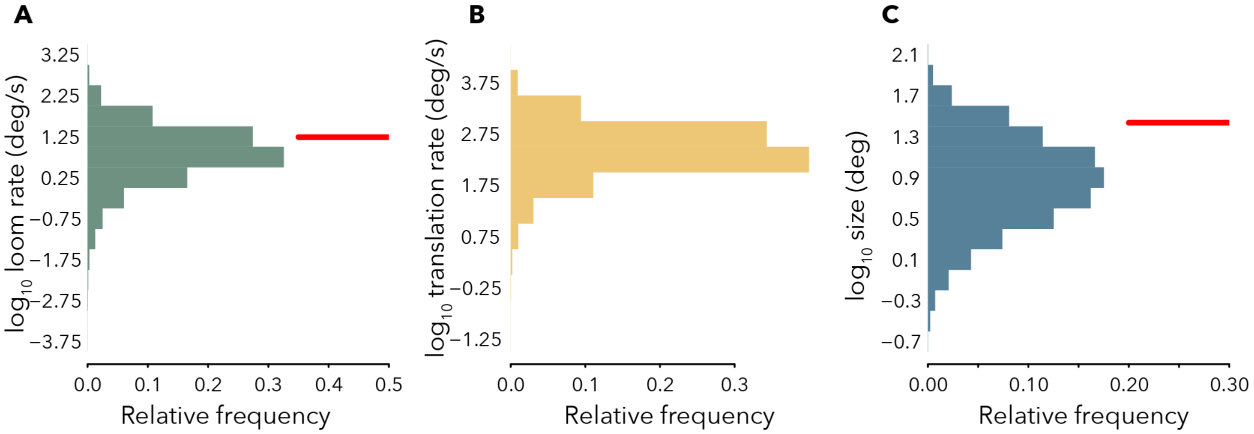


**Figure S4. Visual stimuli produced by neighbors. A.** Instantaneous loom rate, $S'(t)$, produced by neighbors with $S'(t) > 0$ (i.e. those looming rather than receding). Red line shows median of putative lower loom thresholds reported in previous studies of fishes [32]. **B.** Translation rate of neighbors, $T(t)$. **C.** Instantaneous visual size of neighbors, $S(t)$. Red line shows median of putative size thresholds reported in previous studies of fishes [4, 5, 25].

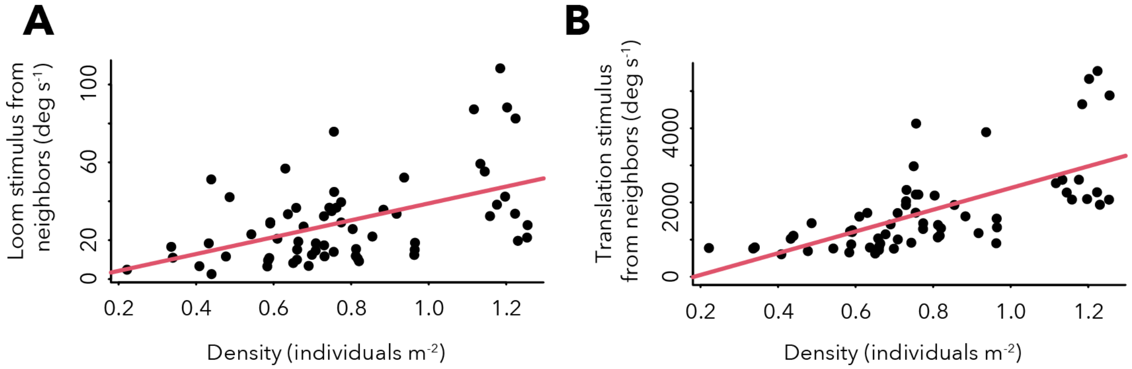


**Figure S5. Sensory stimuli at response time as a function of group density.** Total loom stimulus from neighbors (**A**) computed as the sum of looming stimuli over all visible neighbors (ordinary least squares linear regression: $P = 2.58 \times 10^{-5}$), and total translation stimulus from neighbors (**B**) computed as the sum of translation stimuli over all neighbors for all responding individuals in tracked escape events (ordinary least squares linear regression: $P = 3.02 \times 10^{-9}$). Each point represents visual stimuli perceived by an individual responder. Looming in (A) and translation in (B) were computed over the period starting one second prior to the initiation of the first escape response, and ending at the onset of the escape maneuver by each focal individual. Group density was computed as the average spatial density of individuals in the foraging area over this same period.

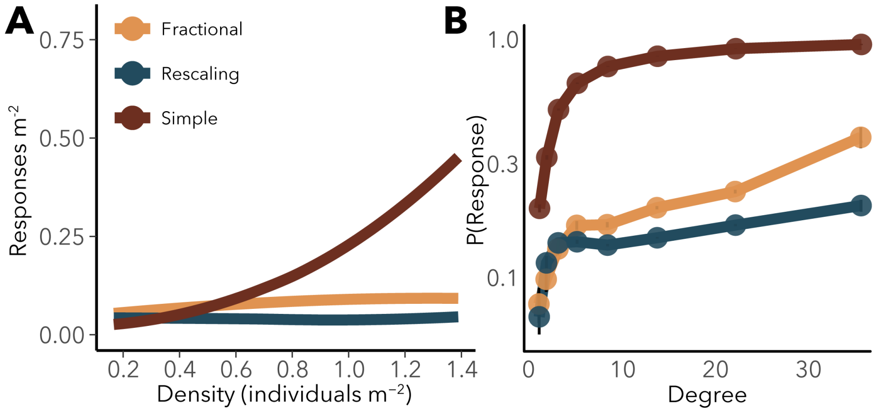


**Figure S6. Exposure to misinformation and local network density. A.** Number of responses as a function of group density. **B.** Scaling of exposure probability, defined as the probability that an individual will see one or more neighbor fleeing over the course of a simulation, as a function of an individual's raw degree in the network (i.e., the number of visible neighbors it has), from simulations in which a misinformation cascade is induced by simulation of an aggressive interaction between a random pair of nearest neighbors in the population (see SI section *Numerical experiment for misinformation spread*).

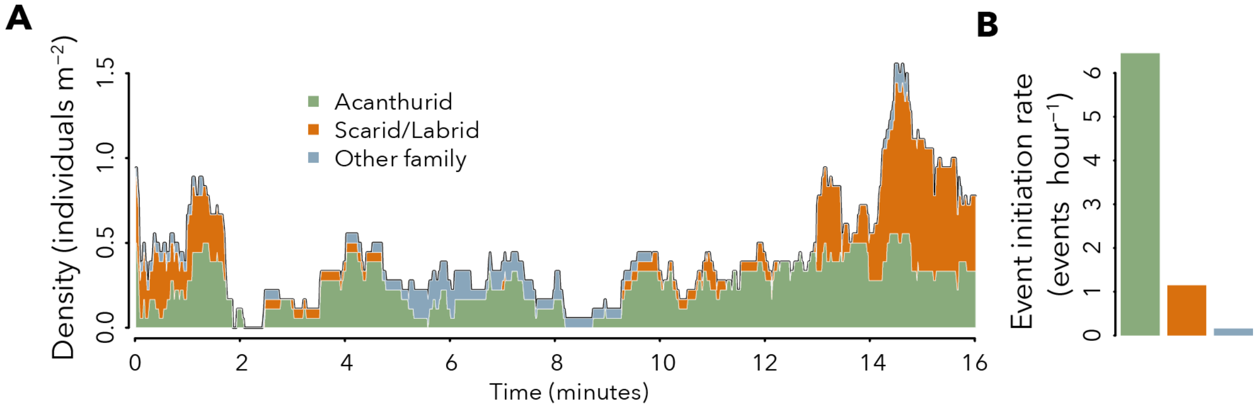


**Figure S7. Fluctuations in density and group compositions. A.** Temporal fluctuations in density and phenotypic (species) composition of fish groups for a representative time series from an unperturbed foraging ground. Colors indicate the density contribution of common reef fish families or family groups: surgeonfishes (Acanthuridae, green), parrotfishes and wrasses (Scaridae/Labridae, orange), and other families (including Zanclidae, Mullidae, Chaetodontidae, and other less common families, blue). **B.** Rate at which each of the most common families/family groups initates escape events computed from 42 escape event time series shown in Fig. 1A of the Main Text. We define an individual as initiator if it was the first responder in a given event.

 320

| Type | Model name | Integration over neighbors | Decision Function / neighbor-wise prob. | Model code |
|---|---|---|---|---|
| Pooling | Linear pooling | $F = \mathbf{V}\boldsymbol{\beta}$ | $\text{logit}(p) = D^{\star} + F$ | p1 |
| | Independent influence | $p = 1 - \prod_i (1 - p_i)^N$ | $F_i = \mathbf{V}_i\boldsymbol{\beta}$, $\text{logit}(p_i) = D^{\star} + F_i$, | p2 |
| | Loom threshold selective attention | $F = \beta_1 S'_{dom}$ | $\text{logit}(p) = D^{\star} + F$ | p3 |
| | Linear selective attention | $F = \mathbf{V}_{dom}\boldsymbol{\beta}$ | $\text{logit}(p) = D^{\star} + F$ | p4 |
| | Tau theory selective attention | $F = \beta_1 S_{dom}[S'_{dom}]^{-1}$ | $\text{logit}(p) = D^{\star} + F$ | p5 |
| | Eta selective attention 1 | $F = \beta_1 S'_{dom} e^{-\beta_2 S_{dom}}$ | $\text{logit}(p) = D^{\star} + F$ | p6 |
| | Eta selective attention 2 | $F = S'_{dom} e^{-\beta_2 \sum_i S_i}$ | $\text{logit}(p) = D^{\star} + F$ | p7 |
| | Loom-gated size selective attention | $F = \beta_1 \mathbf{1}_{[S'_{dom}(t)>0]} S_{dom}$ | $\text{logit}(p) = D^{\star} + F$ | p8 |
| Averaging | Input averaging | $F = \frac{1}{N}\mathbf{V}\boldsymbol{\beta}$ | $\text{logit}(p) = D^{\star} + F$ | a1 |
| | Response averaging | $p = \frac{1}{N}\sum_i^N p_i$ | $F_i = \mathbf{V}_i\boldsymbol{\beta}$, $\text{logit}(p_i) = D^{\star} + F_i$ | a2 |
| Temporal rescaling | Response rescaling | $F = \mathbf{M}\mathbf{w}\,(\gamma + m)^{-1}$ | $\text{logit}(p) = D^{\star} + F$ | rr |

Table 1: Candidate decision-making models describing escape decision-making as a function of incoming sensory stimuli (see *Defining the model set*). Variables, $F(t)$ and $F_i(t)$ denote latent sensory features derived from operations on raw visual sensory input variables, $S(t)$, $S'(t)$, and $T(t)$. Probability, $p(t)$, denotes probability to initiate an escape response in a small time increment, and $p_i(t)$ denotes probability in a small time increment to respond to the $i$th neighbor in the absence of all other neighbors. $N(t)$ is the number of visible neighbors of the focal individual at time, $t$. Vectors, $\boldsymbol{\beta}$ and $\mathbf{w}$, are $(3 \times 1)$ and $(2 \times 1)$ vectors of fitted parameters, respectively. Raw sensory input vectors, $\mathbf{V}(t) := \{\sum_i S_i(t), \sum_i S'_i(t), \sum_i T_i(t)\}$, $\mathbf{V}_\mathbf{i}(t) := \{S_i(t), S'_i(t), T_i(t)\}$, and $\mathbf{V}_{\mathbf{dom}}(t) := \{S_{dom}(t), S'_{dom}(t), T_{dom}(t)\}$, are $(1 \times 3)$ vectors of summed sensory input over all neighbors, input from the $i$th neighbor, and input from the dominant neighbor, respectively. Vector, $\mathbf{M} = \{\sum_i S'_i(t), \sum_i T_i(t)\}$ is the vector of neighbor motion stimuli summed over neighbors. And $D^{\star}$, $\beta_1$, $\beta_2$, and $\gamma$ are scalar fitted parameters. $m(t) = \omega \int_{-\infty}^{t} e^{-\frac{t-s}{\tau}} \mathbf{M}(s)\mathbf{w} ds$ is an exponentially-weighted integral of the past history of sensory input from neighbors with decay timescale, $\tau$, and constant parameter $\omega$. *Integration over neighbors* describes the manner in which information from different neighbors are integrated in the model. For all models except *Independent influence* (p2) and *Response averaging* (a2), integration over neighbors is assumed to occur at the stage of computation of visual features $F$. For *Independent influence* (p2) and *Response averaging* (a2), integration over neighbors is assumed to occur at the stage of response probability computation. Note that time-dependencies of raw visual input variables, $S_i(t)$, $S'_i(t)$, and $T_i(t)$, sensory features, $F(t)$ and $F_i(t)$, number of visible neighbors, $N(t)$, steady state memory, $m(t)$, and response probabilities, $p(t)$ and $p_i(t)$, are omitted to simplify notation.

| Type | Model name | Negative log-Likelihood | Precision-Recall AUC | Class accuracy (FR, NR, SR) |
|---|---|---|---|---|
| Pooling | Linear pooling | 103.08 | 0.52 | 0.70, 0.71, 0.47 |
| | Independent influence | 105.32 | 0.54 | 0.70, 0.61, 0.54 |
| | Loom threshold selective attention | 104.86 | 0.49 | 0.63, 0.62, 0.38 |
| | Linear selective attention | 100.85 | 0.58 | 0.80, 0.53, 0.59 |
| | Tau theory selective attention | 104.36 | 0.47 | 0.77, 0.62, 0.35 |
| | Eta selective attention 1 | 101.09 | 0.61 | 0.80, 0.62, 0.56 |
| | Eta selective attention 2 | 102.50 | 0.55 | 0.77, 0.55, 0.62 |
| | Loom-gated size selective attention | 101.54 | 0.56 | 0.80, 0.56, 0.57 |
| Averaging | Input averaging | 103.63 | 0.53 | 0.70, 0.54, 0.44 |
| | Response averaging | 103.93 | 0.54 | 0.77, 0.54, 0.50 |
| **Temporal rescaling** | **Response rescaling** | **64.28** | **0.86** | **0.80, 0.81, 0.80** |

Table 2: Decision-making model performance. Values in *Class accuracy* column correspond to prediction accuracies for first responders, non-responders, and secodary responders, respectively. Formulation and details of each model are described in Table S1 and *Defining the model set* above.

| Category | Parameter | Interpretation | Units | Value |
|---|---|---|---|---|
| Movement | $w_0$ | Max speed in `foraging` state | body lengths $s^{-1}$ | 1 |
| | $w_1$ | Speed in `flee` state | body lengths $s^{-1}$ | 7 |
| | $u_0$ | Stand. dev. for heading change in `foraging` state | rad | 0.05 |
| | $u_1$ | Stand. dev. for heading change in `flee` state | rad | 0.25 |
| | $d_1$ | Flight duration | s | 0.69 |
| | $r_0$ | Refractory period | s | 0.69 |
| Sensing & decision | $\rho$ | Visual range | m | 2.5 |
| | $\tau$ | Memory decay timescale | s | 0.1 |
| | $D^\star$ | Intercept | nondimensional | -25.0 |
| | $w_{s'}$ | Looming motion gain | $deg^{-1}$ | 7.5 |
| | $w_T$ | Translational motion gain | $deg^{-1}$ | 1.575 |
| | $\gamma$ | Inhibition constant | $s^{-1}$ | 0.022 |

Table 3: Parameter values for individual-based simulations of the response rescaling model.
